# Adult mice with neonatal-like T cell subsets exhibit increased susceptibility to *Bordetella pertussis* and influenza infection

**DOI:** 10.64898/2025.12.11.691875

**Authors:** Matthew R. Hansen, Colleen J. Sedney, Shiyun Xiao, Disha BR Prasad, Wen Zhang, Kalyan Dewan, Jillian Masters, Maiya Callender, Eric T. Harvill, Kimberly D. Klonowski, Nancy R. Manley

## Abstract

Infants are significantly more susceptible to respiratory infection, often resulting in increased morbidity and hospitalization, and occasionally death. This susceptibility is partially explained by the developing nature of the thymus in human infants at, and for several months after, birth. However, the contribution of T cells produced in this thymic microenvironment to infant immune responses has received minimal investigation. Here, we utilized a previously described mouse model (*Foxn1*^Δ*/*Δ^) which exhibits a persistently immature thymus. Through further characterization, we have determined that adult *Foxn1*^Δ*/*Δ^ mice retain some unique T cells observed in neonatal mice including CD8αß^+^ γδ T cells and CD8 T cells displaying a memory-like phenotype. For this reason, we assessed the potential of these neonatal-like T responses to two pathogens which disproportionately affect neonates, *Bordetella pertussis* (*Bp*) and influenza. Utilizing these infections, we demonstrate that T cells generated in an incompletely developed thymus fail to control or mount an effective response against *Bp.* We also observe that *Foxn1*^Δ*/*Δ^ mice control acute influenza infection, a response which does not require IL-17. However, the *Foxn1*^Δ*/*Δ^ mice fail to generate an influenza nucleoprotein (NP) specific CD8^+^ T cell response which is likely associated with their inability to fully clear the infection. Together, these data suggest that *Foxn1*^Δ*/*Δ^ mice can be utilized to study the generation, function, and persistence of some unique T cells made in a neonatal-like thymus.

## Introduction

It is well appreciated that infants are significantly more susceptible to particular infections. These include both viral and bacterial pathogens, such as influenza and *Bordetella pertussis* (*Bp*; whooping cough) [1–3] which combined result in ^∼^190,000-270,000 deaths annually in children under 5 years of age [4]. One of the major differences between neonates and adults are the types of T cells which populate the periphery. Specifically, neonatal CD4^+^ T cells have a general anti-inflammatory or Th2 bias compared to their adult counterparts [5–9]. Neonatal CD8 T cells also favor effector over memory function while some develop a virtual memory (T_VM_) phenotype [5, 10, 11]. Additionally, γδ T cells are some of the earliest T cells to develop and are enriched in neonates compared to adults. These cells often leave the thymus with full effector functions and migrate directly to barrier tissues. The ability to be activated by various stimuli in the absence of TCR engagement has led to T_VM_ and γδ T cells being described as “innate-like”[12–15].

In the context of infection, a lack of pathogen-specific T cells has been associated with infant susceptibility to both *Bp* and influenza [16]. However, new evidence suggests that the neonatal immune system may be fairly effective at combatting infection, at least in the short-term [5, 10, 16]. This is in part due to the development of distinct anti-inflammatory and innate-like lineages that are unique to the neonatal period [5, 6, 9, 14, 17]. How these cells are programmed in the neonatal thymus and exclusively participate in early neonatal immunity is unclear given the rapid transition of mice from a neonatal to juvenile period. The ability to study age-specific T cells by temporally limiting T cell development would assist in answering these questions.

We previously generated a mutation in *Foxn1* (*Foxn1*^Δ*/*Δ^) resulting in a hypomorphic phenotype in which thymus development is arrested at a late fetal stage [18], in contrast to *Foxn1^nu/nu^* nude mice which lack all thymic development. These *Foxn1*^Δ*/*Δ^ mutants may provide a model with temporally limited T cell output to the late fetal period [18–22]. Further analysis of the T cells generated in these mice confirmed a high frequency of unconventional, innate-like virtual memory T cells (T_VM_) and identified CD8αß^+^ γδ T cells in the periphery of naïve adult *Foxn1*^Δ*/*Δ^mice. *Foxn1*^Δ/Δ^ mice were highly susceptible to *Bp*, associated with a failure to generate a protective CD4^+^ T cell response. In comparison, inoculation of *Foxn1*^Δ/Δ^ mice with influenza resulted in a unique response with an initial reduction in viral titers, and weight recovery akin to control mice; however, these *Foxn1*^Δ/Δ^ mice ultimately failed to clear the infection. The acute response was not significantly dependent on IL-17 but associated with a prepositioning of the T_VM_ and CD8αß^+^ γδ T cells in the lung parenchyma. However, failure to clear influenza infection was associated with the lack of an antigen-specific CD8^+^ T cell response. This model provides novel observations on unconventional T cell responses to respiratory pathogens, and future possibilities for studying the underlying mechanisms that support the development of certain T cells unique to the late fetal thymus.

## Methods

### Mice

Age matched, 8-12 week old, and sex matched *Foxn1*^Δ*/*Δ^ and littermate *Foxn1^+/^*^Δ^ mice were generated by Dr. Manley’s laboratory as previously described [18], bred, and maintained in-house. *IL17af^-/-^* mice were ordered from the JAX lab (strain #: 034140). *IL17af^-/-^* were crossed with *Foxn1*^Δ*/*Δ^ to generate *IL17af^-/-^* mice on the *Foxn1*^Δ*/*Δ^ background (*Foxn1*^Δ*/*Δ^, *IL17af^-/-^)* with (*Foxn1*^Δ*/*Δ^*; IL-17af^+/-^*) as controls. The C57Bl/6 mice used for the neonatal studies were bred in-house and originally obtained from Charles River Laboratories. All mice were maintained in a pathogen-free facility at the University of Georgia. All animal experiments were approved by the University of Georgia Institutional Animal Care and Use Committee.

### Pathogens and Infections

The *B. pertussis* strain Tohama 1 (WT *B. pertussis*) was prepared as previously described [23]. Bacteria were maintained on Bordet-Gengou agar (BD) supplemented with 10% defibrinated sheep blood (Hemostat) and 20 mg/ml gentamicin. Liquid cultures were grown overnight in Stainer-Scholte broth at 37° C to mid-log phase then maintained in 20% glycerol stocks at -80° C. Adult female and male mice were infected intranasally (i.n.) with 5 × 10^5^ CFU *Bp* /50 µl of PBS. The influenza A virus A/HK-x31 (x31, H3N2) was generously provided by Dr. S. Mark Tompkins (University of Georgia, Athens, GA). Adult female and male mice were i.n. infected with 10^3^ PFU of x31in 50ul of PBS. At the indicated timepoints, mice were euthanized via CO_2_ inhalation (Bp) or anesthetized with 2% 2,2,2 tribromoethanol solution (influenza) delivered intraperitoneally followed by cervical dislocation and tissue extraction.

### Tissue extraction and flow cytometry

For *Bp* infections, lungs were processed as previously described [23]; for influenza, single cell suspensions from tissues were obtained as previously described [24]. Bronchoalveolar lavage (BAL) was collected by inserting a catheter into a small incision in the trachea and flushing the lungs with three 1 mL volumes of PBS. Lungs were excised, minced, and incubated for 30 min at 37°C with 1.25mM EDTA followed by a 1 h incubation with 150 units/mL collagenase (Life Technologies, Grand Island, NY). Cells were then passed through a 40 μM cell strainer and resuspended in 44% Percoll underlaid with 67% Percoll, centrifuged, and the cellular interface collected. Lymph nodes and spleens were mechanically processed and passed through cell strainers. Erythrocytes were removed using Tris-buffered ammonium chloride. For the intravascular (I.V.) staining, animals received 3ug of biotinylated αCD45 (clone 30-F11) mAb delivered via the tail vein 3 min prior to euthanasia. For phenotypic analysis of various lymphoid populations, cells were surface stained using various combinations of fluorescently conjugated antibodies as described of the indicated reactivity (clone) for 20 minutes at 4°C: FITC-conjugated αTCR γδ^+^ (GL3), αIFNγ (XMG1.2), αCD44 (IM7), αCD3 (17A2), or αCD8α^+^ (2.43); PercpCy5.5-conjugated αCD4^+^ (RM4-5), αTCRß (H57-597), or αCD44 (IM7); APC-conjugated αCD8ß (YTS156.7.7), αEomes (Dan11mag); VioletFluor450-conjugated αCD8α^+^ (2.43); Brilliant Violet 650 conjugated αCD8α^+^ (53-6.7), PE-conjugated αCD8α^+^(2.43); PE-Cy7-conjugated αCD8α^+^ (2.43), αCD44 (IM7), or CD4^+^ (RM4-5); Alexafluor 700 conjugated αCD4^+^ (RM4-5) APC-Fire 750 conjugated αCD49d (R1-2); PE/Cy5 conjugated αTCR γδ^+^(GL-3); Antibodies purchased from BD Biosciences (Franklin Lakes, NJ), Biolegend (San Diego, CA), Tonbo (Tucson, AZ), or Invitrogen (Waltham, MA). Intracellular staining for Eomes was performed as previously described [25]. Where Zombie Aqua fixable viability dye (Biolegend, San Diego, CA) was used, cells were stained for viability prior to surface staining with fluorophore conjugated mAbs, according to manufacturer’s protocol. For influenza antigen-specific responses: influenza nucleoprotein (NP) MHC class I tetramer [H-2D(b)/ASNENMETM; NP_366-374_] conjugated to Brilliant Violet 421 was generated at the National Institutes of Health Tetramer Core Facility (Emory University, Atlanta, GA). Staining with NP-tetramer was carried out at RT for 1 h in conjugation with other surface staining mAbs. Intravascular stained samples were incubated with streptavidin-conjugated Brilliant Violet 421 for 15 min at 4°C followed by surface staining with mAbs. All samples were fixed with 2% paraformaldehyde or 20 min at 4°C prior to flow cytometry. Data was acquired using a NovoCyte Quanteon with NovoExpress Software (Agilent, Santa Clara, CA) and analyzed using FlowJo software version 10.10.0 (BD Biosciences Franklin Lakes, NJ). All samples were gated on single cells and lymphocytes prior to further analysis.

### In vitro T cell stimulation

Bulk splenocytes or bulk leukocytes from the lung were cultured in the presence of 5 ng/mL PMA and 500 ng/mL Ionomycin, αCD3/αCD28 coated wells, or a cocktail of 10 ng/mL IL-12, 10 ng/mL of IL-18, and 100 U/mL IL-2 (eBioscience, San Diego, CA) in RPMI with 10% FCS and 10% supplementum completum (penicillin-streptomycin, L-glutamate, HEPES, B-mercaptoethanol, gentamycin, and FCS) at 37°C. Cells were stimulated in the presence of brefeldin A (BFA) for 3hrs (PMA/ Ionomycin), 5h (αCD3/αCD28 stimulation), or 5h (after IL-12/Il-18/IL-2 stimulation for 18h without BFA). Cells were subsequently surface stained and fixed overnight in 2% PFA. The following day, cells were permeabilized using eBioscience Perm/Wash and intracellularly stained per the manufacturer (eBioscience, San Diego, CA) for IFNγ (XMG1.2), IL-17a (TC11-18H10.1), and IL-4 (11B11). Lastly, cells were fixed in 2% PFA for FACS analysis.

For assessment of intracellular cytokine production in response to an influenza-specific epitope, cells were harvested and re-stimulated with either 1 µg/mL of the influenza A NP_366-374_ peptide or a non-specific peptide [SIINFEKL] diluted in RPMI with 10% FCS, 10% supplementum completum and golgistop (BFA) for 5 hours at 37°C. Following re-stimulation, cells were stained for cell surface markers and fixed overnight in 3% PFA. Cells were then permeabilized using eBioscience Perm/Wash, stained, and resuspended in 2% PFA for flow cytometry.

### Assessment of bacterial burden

For determining Bp bacterial burden, organs were excised and homogenized in 1 ml PBS, serially diluted, and plated on Bordet Gengou agar (BD) supplemented with 10% defibrinated sheep blood (Hemostat). Colonies were counted following a 5 d incubation at 37° C to quantify bacterial numbers.

### IFNγ ELISA

Lung homogenate supernatant samples were collected and stored at -20C prior to cytokine analysis. Samples were assessed for concentrations of IFNγ utilizing the R&D Systems DuoSet ELISA kit (Minneapolis, MN) per the manufacturer’s instructions.

### Influenza plaque assay

Lung samples from dissected mice were suspended in 1 ml PBS and mechanically homogenized for 60s using a Tissue-Lyzer II (Qiagen, Hilden, Germany), followed by centrifugation at 7,000g for 10 mins. The supernatant was collected and stored in -80° C for use in plaque assays. As previously described [26], MDCK cells were cultured in DMEM containing 10% Fetal Bovine Serum (Gibco, Waltham, MA) in 12-well plates. Lung homogenate samples were serially diluted in an infection medium containing 1ug/µl TPCK-trypsin. These dilutions were introduced to the cells in duplicates and the plates were incubated at 37°C for 1 h, with gentle rocking every 15 min to ensure uniform distribution of the infection medium. The cells were incubated with an overlay medium containing 1.2% Avicel microcrystalline cellulose (FMC BioPolymer, Philadelphia, PA) for 72 h. The overlay was removed, and the cells were subsequently washed, fixed with cold methanol/acetone (60:40%), and stained with crystal violet.

### Lung injury assay

Lung injury assay was performed as previously described [27]. Briefly, 200 µL of 0.5% Evan’s blue dye (Fisher Scientific, Waltham, MA) diluted in PBS was injected intravenously via the tail vein. After 1 h, BAL was collected and lungs perfused with PBS to remove any Evan’s blue dye in the vasculature. For lung samples, Evan’s blue dye was extracted in formamide (Sigma Aldrich, Burlington, MA) and quantified as absorbance at 620 nm using a multi-modal microplate reader (Biotek, Santa Clara, CA).

### Statistical Analysis

For experiments comparing two groups, an unpaired two-tail Student’s T test was performed. Where three groups were compared, a one-way ANOVA with Tukey’ test was performed. For experiments involving 3 groups and 2 stimulation conditions, a two-way ANOVA with Tukey’s test was performed. Statistical calculations were performed using GraphPad Prism 10.0.3 for Windows (Graphpad Software, Boston, MA).

## Results

### Phenotypic characterization of T cells in lungs of *Foxn1*^Δ/Δ^ mice

*Foxn1*^Δ/Δ^ mice were previously generated via N-terminal deletion of the *Foxn1* gene [18]. These mice were found to harbor a reduced number of circulating and secondary lymphoid organ residing T cells and, of the T cells present, a majority display a memory-like phenotype (CD44^hi^ CD62L^lo^) [20, 21]. As thymus development appears to arrest at a late fetal stage in these mice, we hypothesized that as adults, they may harbor neonatal-like T cells and thus could provide a unique opportunity to evaluate the function of these cells in the context of infection. Given the memory-like phenotype of the T cells in the *Foxn1*^Δ/Δ^ mice, we first sought to determine whether the T cells in these mice are predisposed to populate the respiratory tract.

We first performed intravascular staining by delivering an αCD45 mAb I.V. 3 min prior to animal euthanasia to distinguish populations of circulating (CD45 I.V.^+^) and tissue-embedded (CD45 I.V.^-^) cells. Significantly, *Foxn1*^Δ/Δ^ mice have approximately 13x the frequency of CD4^+^ and 8x the frequency of CD8^+^ T cells in the lung parenchyma (CD45 I.V.^-^) compared to *Foxn1*^+/Δ^ mice (**Figure 1A-B**). However, taking the lower numbers of circulating T cells in *Foxn1*^Δ*/*Δ^mice into consideration, the absolute count of CD45 I.V.^-^ T cells is ∼10x more CD4^+^ and ∼5x more CD8^+^ T cells in the lung parenchyma (CD45 I.V.^-^) of *Foxn1*^Δ*/*Δ^mice, and remains significant. Therefore, it is likely that the high CD44 expression and low CD62L expression on T cells in the *Foxn1*^Δ/Δ^ mice [20] facilitated the trafficking of these cells into non-lymphoid tissues. In adults, non-lymphoid tissues are populated by memory cells generated from past infections that maintain a frontline barrier to secondary infection [28, 29]. However, previous studies have identified memory-like T cells in the periphery of naïve neonatal mice, collectively referred to as virtual memory T cells (T_VM_) [10]. T_VM_ are predominately CD8^+^ T cells that express conventional memory cell markers (CD44^hi^CD122^+^), among others (CD49d^lo^Eomes^+^) [10, 30] without any prior immunization. Since the *Foxn1*^Δ/Δ^ mice harbored a greater frequency of CD8^+^ T cells in their lung parenchyma compared to *Foxn1*^+/Δ^ littermates, we hypothesized that many of these cells may be T_VM_. To determine whether T_VM_ cells populate the respiratory tract of *Foxn1*^Δ/Δ^ mice, we assessed the frequency and number of T_VM_ (CD8^+^CD44^hi^CD122^+^) in the lung. Indeed, *Foxn1*^Δ/Δ^ mice harbored a significantly greater proportion (∼2x) of T_VM_ in their lungs compared to *Foxn1*^+/Δ^ mice, while absolute counts were similar between *Foxn1*^Δ/Δ^ mice and *Foxn1*^+/Δ^ mice (**Figure 1 C-D**). In the spleen, we also observed a significant increase in the proportion of T_VM_ (∼2x) in *Foxn1*^Δ/Δ^ mice, but a decrease in absolute T_VM_ number (**Figure F-G**). Interestingly, the majority of T_VM_ from *Foxn1*^Δ/Δ^ lungs expressed high levels of CD49d compared to T_VM_ cells from *Foxn1*^+/Δ^ mice, where the majority express low levels of CD49d (**Figure 1E**). This observation was absent in T_VM_ isolated from the spleens of *Foxn1*^Δ/Δ^ mice, which were phenotypically similar to *Foxn1*^+/Δ^ mice (CD49d^lo^ Eomes^+^) (**Figure 1H**).

**Figure 1:**
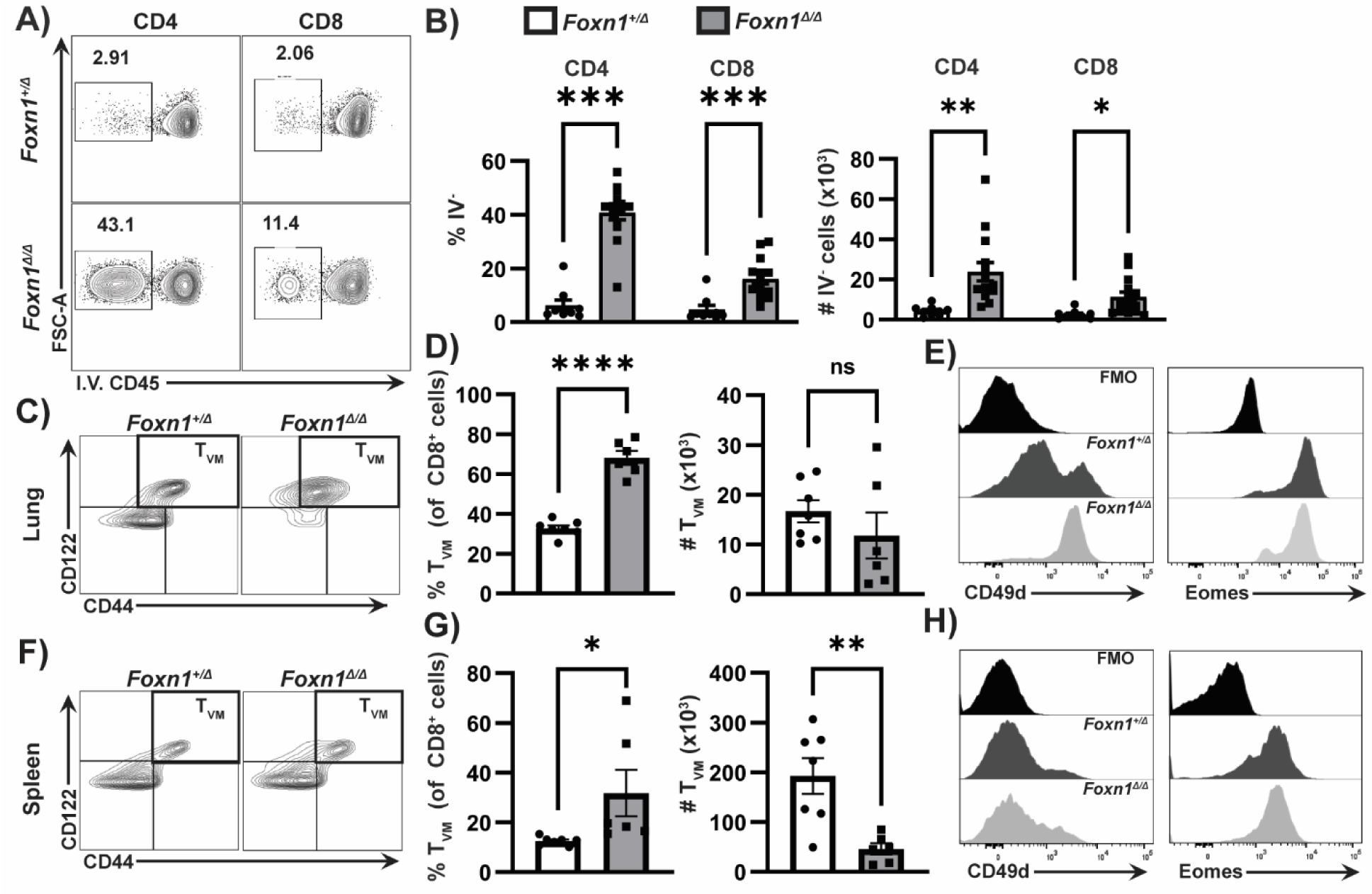
*Foxn1*^Δ*/*Δ^mice harbor increased numbers of lung-embedded T cells with an increased frequency of virtual memory T cells (T_VM_). (A) Representative flow cytometry plots of lung-embedded (CD45 I.V.^-^) CD4^+^ and CD8^+^ T cells labelled by I.V. administration of αCD45; previously gated on lymphocytes, single cells, TCRß^+^ and CD4^+^ or CD8α^+^cells. B) Frequency (left) and total cell number (right) of I.V.^-^ CD4^+^ and CD8^+^ T cells in *Foxn1^+/^*^Δ^ and *Foxn1*^Δ*/*Δ^lungs (n=9-14 mice per group). Representative plots of T_VM_ (CD44^hi^CD122^+^) within the CD8α^+^ gate (not shown) in *Foxn1^+/^*^Δ^and *Foxn1*^Δ*/*Δ^ lungs (C) and spleen (F). Frequency (left) and total cell numbers (right) in *Foxn1^+/^*^Δ^and *Foxn1*^Δ*/*Δ^ lungs (D) and spleen (G) (n=6-7 mice per group). Representative plots of expression of CD49d (left) and Eomes (right) amongst T_VM_ from *Foxn1^+/^*^Δ^and *Foxn1*^Δ*/*Δ^ lungs (E) and spleen (H). Error bars show standard error of the mean. All experiments were completed in duplicate. Statistical significance was calculated via unpaired two-tailed T test. (*p=<0.05, **=p<0.01, ***=p<0.001).

γδ T cells also seed barrier tissues early in life and like T_VM_, may also be poised in the lung of *Foxn1*^Δ/Δ^ mice to respond to infection. Therefore, we assessed the frequency of various phenotypes of γδ T cells (**Figure 2A-C)** within the CD45 I.V.^-^ pool. The frequency of CD4^-^CD8^-^ (DN) γδ T cells embedded in the lung tissue was similar between *Foxn1*^Δ/Δ^ mutant and *Foxn1*^+/Δ^ control mice (**Figure 2B-C)**. While CD8αα^+^ γδ T cells were similar in frequency in the lung tissue of both animal strains, they were increased numerically in the *Foxn1*^Δ/Δ^ mice (**Figure 2B-C)**. Interestingly, there was a significant increase in the frequency of CD45 I.V.^-^ γδ T cells in the lungs of *Foxn1*^Δ/Δ^ mice that expressed heterodimers of CD8α and CD8ß (CD8αß^+^), ∼19% versus 9% in *Foxn1*^+/Δ^ mice (**Figure 2A-C**). While CD8αα^+^ γδ T cells are well described as intraepithelial lymphocytes (IELs) [31–33], the observation of a significant number of CD8αß^+^ γδ T cells in *Foxn1*^Δ/Δ^ lungs (**Figure 2A-C**) was surprising. γδ T cells from adult WT mice are not typically reported to express CD8αß heterodimers; however, these cells could be unique to the fetal thymus and neonatal period. Indeed, an average of 25% of the T cells (CD3^+^) recovered from the lung of newborn C57BL/6 mice (P0) were CD8αß^+^ γδ T cells; however, these cells wane rapidly by post-natal day 7 (P7) (**Figure 1D, E**). Recent work from the Pennington lab demonstrated that CD8αß^+^ γδ T cells develop perinatally [17], which aligns with our data in *Foxn1*^Δ/Δ^ and newborn wild-type mice (**Figure 2A-E**). In summary, these results suggest that *Foxn1*^Δ/Δ^ mice are enriched for unique neonatal-like T cell populations (T_VM_ and CD8αß^+^ γδ T cells) in the lung.

**Figure 2:**
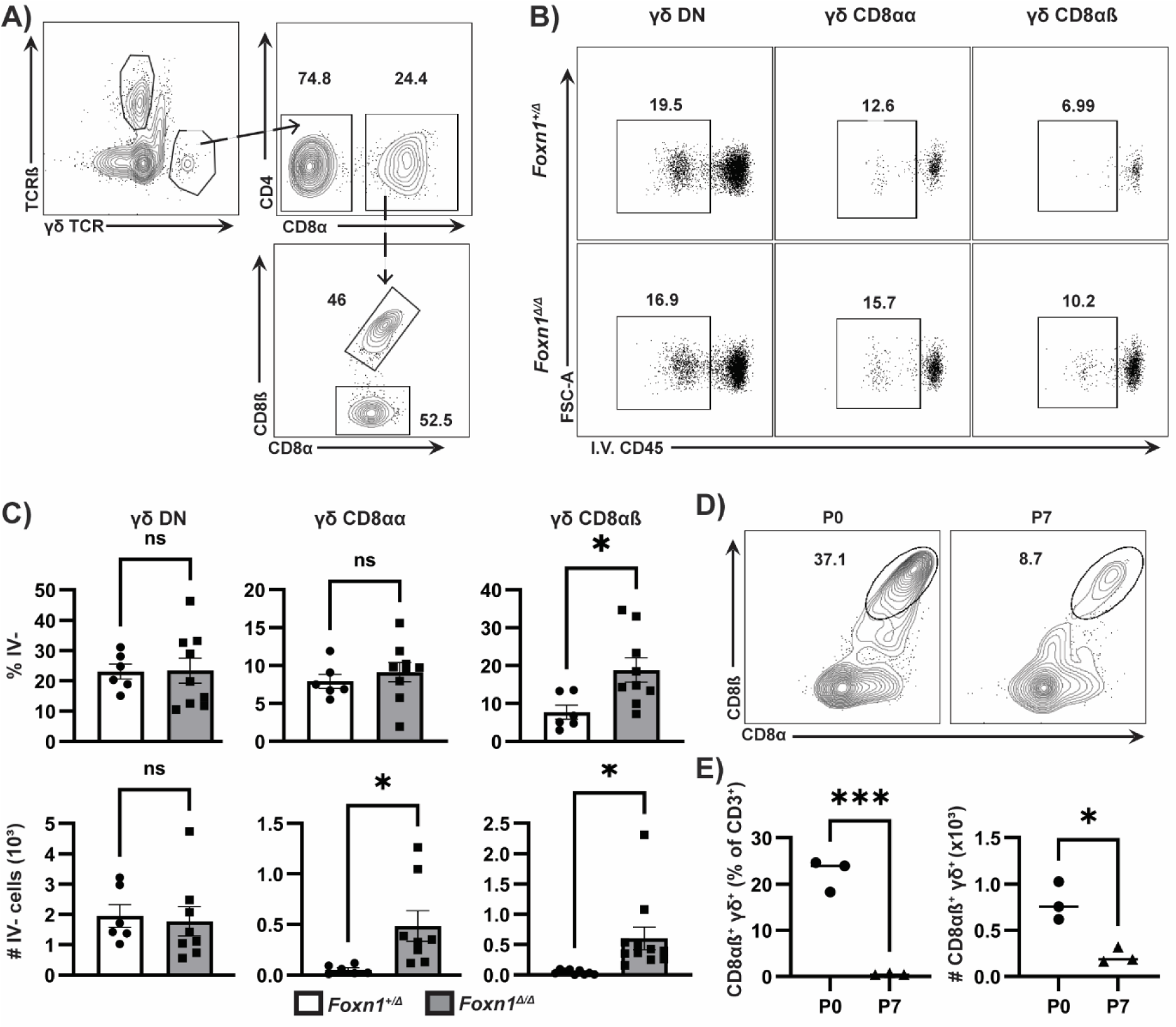
Populations of γδ^+^ T cells enriched in the lung of *Foxn1*^Δ*/*Δ^ mice. (A) Representative gating strategy of γδ T cell subsets: DN (CD4^-^CD8α^-^) within the γδ^+^ cells, and CD8αα^+^ and CD8αß^+^ γδ T within the CD8α^+^ γδ T cell gate; previously gated on lymphocytes and single cells (not shown). (B) Representative flow cytometry plots of CD45 I.V.^-^ cells among γδ T cell subsets defined by the gating strategy in panel A. (C) Frequency (top) and total cell number (bottom) of lung embedded (CD45 I.V.^-^) γδ T cells based on subset (n=6-8 mice per group). (D) Representative flow cytometry plots of DN, CD8αα^+^, and CD8αß^+^ γδ T cells in newborn (P0) and P7 C57BL/6 pup lungs; previously gated on CD3^+^ lymphocytes and single cells. (E) Total cell number (right) and frequency within the CD3^+^ population (left) of CD8αß^+^ γδ T cells in the lungs of C57BL/6 pups at P0 and P7. Error bars show standard error of the mean. Experiments were completed in duplicate. Statistical significance was calculated via unpaired two-tailed T test. (*p=<0.05, **=p<0.01, ***=p<0.001).

### Functional characterization of T cells in *Foxn1*^Δ/Δ^ mice

T_VM_ and γδ^+^ T cells can be activated in a TCR dependent or independent fashion, the latter also referred to as bystander activation. Activated T_VM_ and γδ^+^T cells can exhibit a variety of effector functions including cytotoxic killing and production of effector cytokines. Infection of mucosal sites leads to a rapid burst of IFNγ, often provided by bona-fide tissue resident memory cells, that stimulates tissue macrophages and dendritic cells, and potentiates a Th1 adaptive response [34, 35]. We hypothesize that T_VM_ and γδ T cells positioned within the lung of *Foxn1*^Δ*/*Δ^ are poised to respond to pathogens in this manner. To begin to explore this possibility, we sought to assess the functional capacity of these lung embedded innate-like cells and compare their response to cells isolated from both *Foxn1^+/^*^Δ^ controls and P5 C57BL/6 neonates.

We performed intravascular staining using a biotin-conjugated αCD45 mAb on *Foxn1*^Δ*/*Δ^ and *Foxn1^+/^*^Δ^ mice or perfused the lungs of the P5 C57BL/6 neonates, and subsequently stimulated the isolated cells with PMA/Ionomycin. In addition to assessing IFNγ, we also measured IL-17a due to the important role of this cytokine in T cell responses to extracellular bacteria like *Bp* [36, 37]. Significantly, ∼70% of the CD45 I.V.^-^ negative or lung tissue embedded CD8^+^ T cells produced IFNγ in *Foxn1*^Δ*/*Δ^ mice, ∼4-fold higher than *Foxn1*^+/Δ^ control mice and the P5 neonate (**Figure 3A, B**). Among the CD45 I.V.^-^ CD4^+^ cells in the lung, IFNγ production was also significantly higher in *Foxn1*^Δ/Δ^ mice (∼35%) compared to control mice (∼13%), and much lower in the P5 neonate (1%) (**Figure 3 C**). Interestingly, the I.V.^-^ CD4^+^ cells from *Foxn1*^Δ/Δ^ mice readily produced IL17a after PMA/Ionomycin stimulation at ∼7x higher frequency compared to *Foxn1*^+/Δ^ controls and P5 neonates (**Figure 3D**). The increased quantity of lung-embedded CD4^+^ T cells (**Figure 1A, B**), combined with their greater production of either IFNγ or IL17a production (**Figure 3C, D**), suggests that *Foxn1*^Δ/Δ^ mice harbor a relatively large pool of hyperresponsive CD4^+^ T cells in the tissue poised for effector function. We next assessed the cytokine production by CD45 I.V.^-^ γδ^+^ T cells and did not observe a significant difference in either IFNγ or IL-17a production after PMA/ Ionomycin stimulation between the groups of mice (**Figure 3E, F**). While we were particularly interested in the behavior of CD8αß^+^ γδ T cells, the ability to reliably capture enough of the different γδ^+^ T cell phenotypes after lung tissue processing and subsequent culture was challenging. For this reason, we utilized splenocytes which also harbored a significant population of CD8αß^+^ γδ T cells (**Supplementary Figure 2A**).

**Figure 3:**
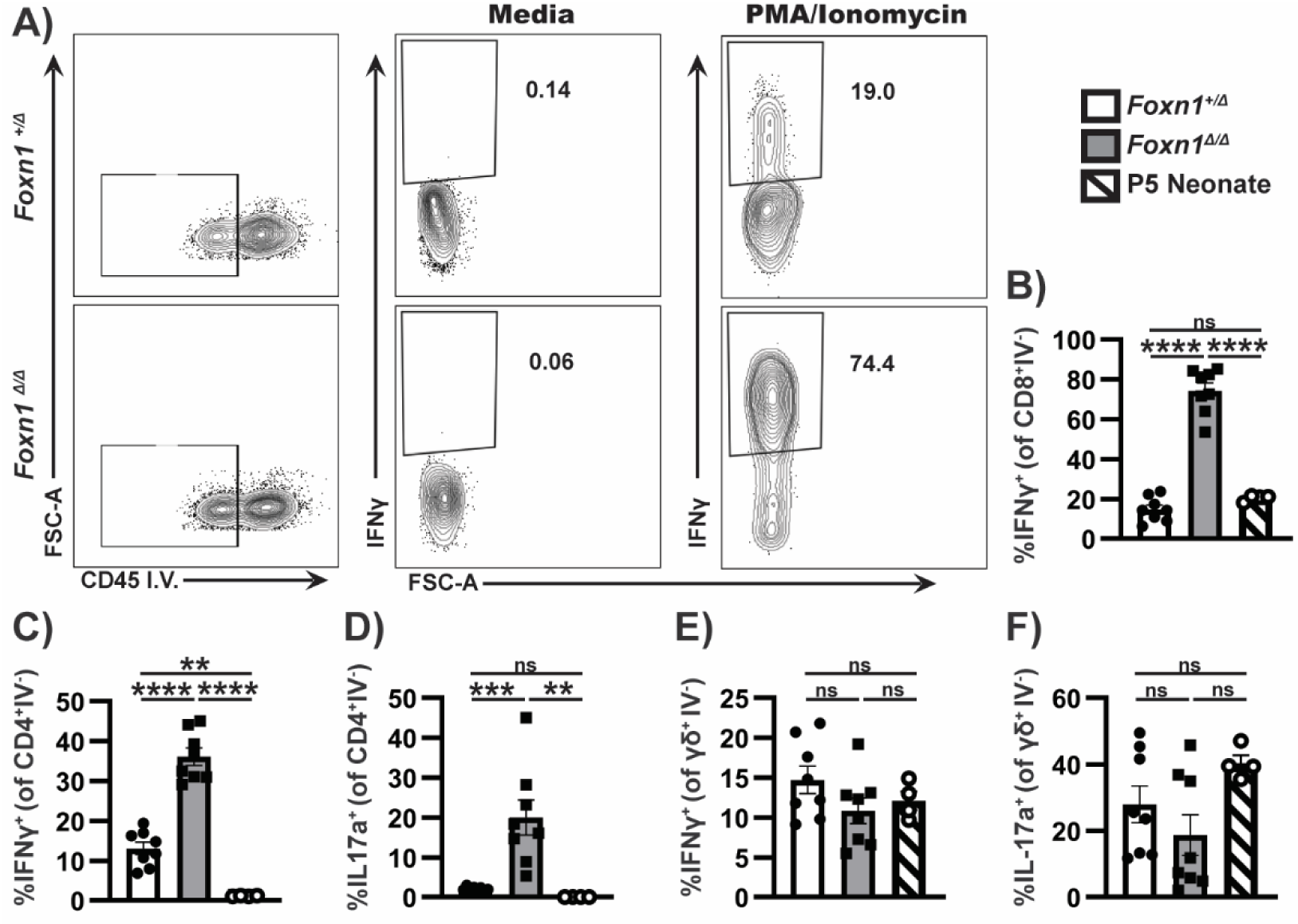
Functional characterization of *Foxn1*^Δ*/*Δ^ versus *Foxn1*^+/Δ^ and neonatal lung embedded T cells. (A) Representative flow cytometry plots and quantification (B) of IFNγ expression after stimulation with media alone (middle) or with PMA/Ionomycin (right); previously gated on CD45 I.V.^-^ cells (left), *Foxn1*^Δ/Δ^ (white) and *Foxn1*^Δ/+^ (grey) only, and TCRß^+^ CD8^+^ lymphocytes and single cells (not shown); P5 neonates (stripes) were not I.V. stained but isolated from a perfused lung. Quantification of IL-17a or IFNγ expression by lung T cells after stimulation with PMA/Ionomycin; previously gated on single cell lymphocytes, CD45 I.V.^-^ (*Foxn1*^Δ/Δ^ and *Foxn1*^Δ/+^ only) TCRß^+^ CD4^+^ (C-D) or TCRß^+^ CD8α^+^ (B) or γδ^+^ (E-F) cells. Experiments were completed in duplicate. Error bars show standard error of the mean (n=8 mice per group). Statistical significance was calculated via a one-way ANOVA with Tukey’ test. (*p=<0.05, **=p<0.01, ***=p<0.001).

After confirming the presence of phenotypically similar splenic γδ^+^ T cells in *Foxn1*^Δ/Δ^ mice (**Figure 4A**), we assessed the ability of T cells isolated from the spleen to respond to both TCR mediated and bystander cytokine mediated activation. Splenocytes were stimulated with αCD3/CD28 or a cocktail of cytokines known to support bystander activation: IL-2/12/18. γδ T cells display heterogeneity in phenotype and function, including distinct subsets of IFNγ and IL-17a producing cells [14], and we also observed a diversity in response to various stimuli. DN γδ T cells from *Foxn1*^Δ/Δ^ mice and P5 neonates showed a significant 4-fold increase in the frequency of IFNγ production after αCD3/CD28 stimulation compared to *Foxn1*^+/Δ^ controls (**Figure 4C**). The response of DN γδ T cells to bystander activation was similar between *Foxn1*^Δ/Δ^ and *Foxn1*^+/Δ^ controls, with roughly 30% IFNγ^+^ cells, which was roughly half the frequency observed in the P5 neonate (∼60%.) (**Figure 4C)**. IL-17a production by DN γδ T cells was also similar in *Foxn1*^Δ/Δ^ and *Foxn1*^+/Δ^ mice (∼3.5%), which was somewhat higher than what was observed in the neonate but only statistically significant when compared *Foxn1*^+/Δ^ (**Figure 4E**). CD8αα^+^ γδ T cells are known IFNγ producers [38], and irrespective of the stimuli, these T cells produce IFNγ similarly in *Foxn1*^Δ/Δ^ and *Foxn1*^+/Δ^ mice (**Figure 4B**). We did not recover a significant number of CD8αα^+^ γδ cells from the lungs of P5 neonates and thus could not analyze IFNγ production for these cells in this group of mice. Significantly, a high frequency of CD8αβ^+^ γδ T cells from *Foxn1*^Δ/Δ^ mice and the P5 neonate expressed IFNγ after cytokine stimulation (∼60%); production of IFNγ was also significantly increased in the same cell population after αCD3/CD28 stimulation (∼30%) (**Figure 4D**).

**Figure 4:**
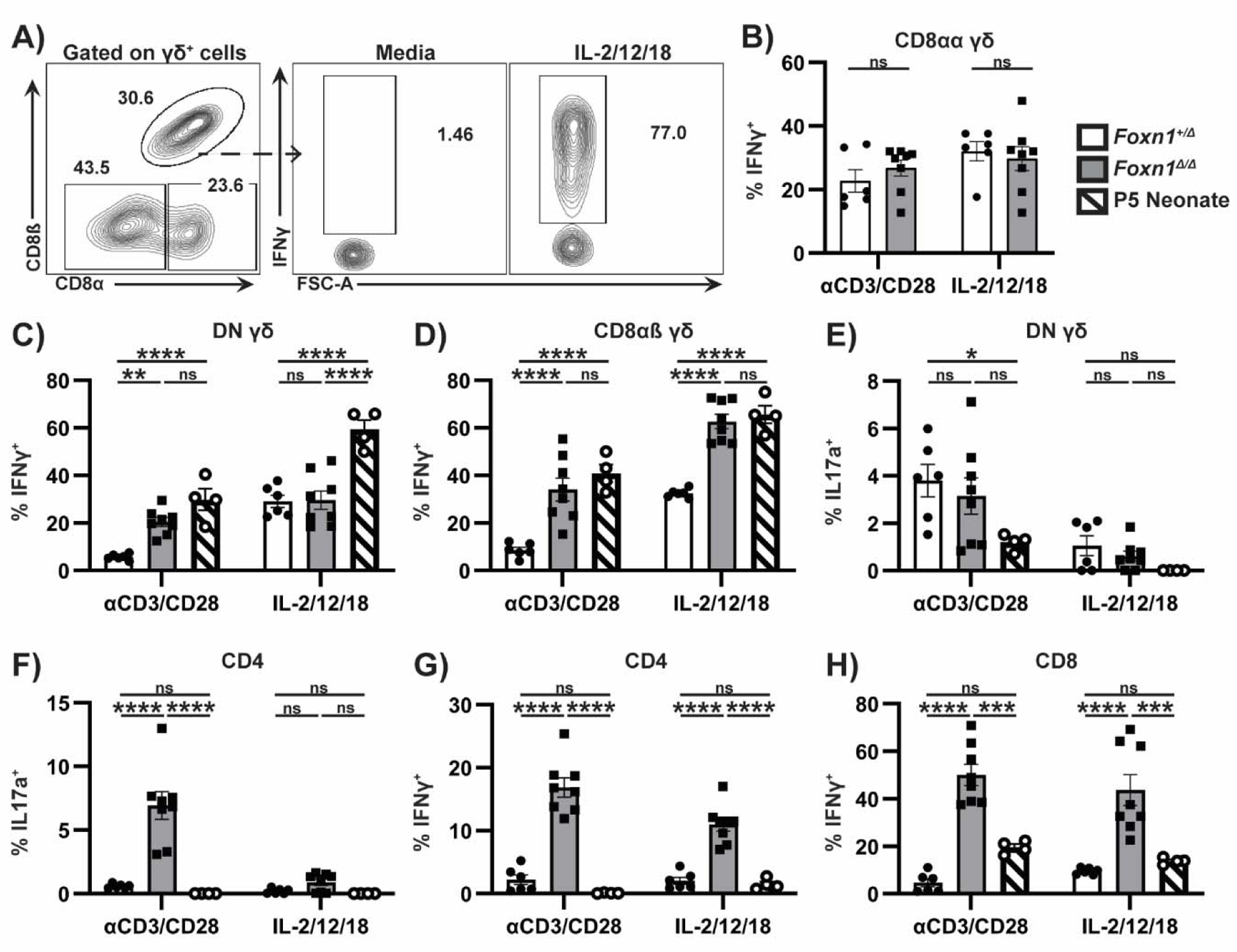
T cells from *Foxn1*^Δ/Δ^ mice are hyperresponsive to TCR mediated and bystander activation. (A) Representative gating of IFNγ^+^ *Foxn1*^Δ*/*Δ^ splenocytes after stimulation with media alone or including the cytokines IL-2, IL-12, IL-18 (middle and right); previously gated on single cell lymphocytes, TCR γδ^+^ (not shown) followed by CD8α^+^ CD8ß^+^ (left). Frequency of *Foxn1^+/^*^Δ^ (white), *Foxn1*^Δ*/*Δ^ (grey), and P5 neonate (striped) CD8αα^+^(B), DN (CD4^-^CD8^-^) (C), and CD8αß^+^ (D) γδ^+^ T cells producing IFNγ in response to αCD3/αCD28 or cytokines. (E) Frequency of DN (CD4^-^CD8^-^) γδ^+^ T cells producing IL17a in the indicated mouse strains after αCD3/αCD28 and cytokine stimulation. Frequency of *Foxn1^+/^*^Δ^, *Foxn1*^Δ*/*Δ^, and P5 neonate CD4^+^ (TCRß^+^) T cells producing IL17a (F) or IFNγ (G) in response to the indicated stimuli. (H) % IFNγ^+^of *Foxn1^+/^*^Δ^, *Foxn1*^Δ*/*Δ^, and P5 neonate CD8^+^ (TCRß^+^) T cells in response to αCD3/αCD28 (left) and cytokines (right). Experiments completed in duplicate (n=6-8 mice per group). Error bars show standard error of the mean. Statistical significance was calculated via two-way ANOVA with Tukey’s test. (*p=<0.05, **=p<0.01, ***=p<0.001).

Similar to PMA/Ionomycin stimulation of lung embedded CD4^+^ cells (**Figure 3D**), a significantly higher portion (∼25%) of splenic CD4^+^ T cells from *Foxn1*^Δ/Δ^ mice produced IL-17a in response to αCD3/CD28 stimulation (**Figure 4F**), 10x more than in *Foxn1*^+/Δ^ mice and P5 neonates. The frequency of IFNγ production in *Foxn1*^Δ/Δ^ CD4^+^ cells was approximately 8-fold and 6-fold higher than *Foxn1*^+/Δ^ mice and P5 neonates in response to αCD3/CD28 and bystander activation, respectively (**Figure 4G**). In response to either stimulus, *Foxn1*^Δ/Δ^ splenic CD8^+^ T cells produced IFNγ at a statistically higher frequency compared to those same cells in *Foxn1*^+/Δ^ controls and P5 neonates, ∼10x and 5x higher for αCD3/ CD28 and the cytokine cocktail, respectively (**Figure 4H**). This suggests that CD8^+^ T cells in *Foxn1*^Δ/Δ^ mice are more sensitive to these activating signals. In summary, *Foxn1*^Δ/Δ^ mice harbor peripheral CD4^+^ and CD8^+^ T cells that are readily activated by a variety of stimuli to produce IFNγ. Importantly, CD4^+^ T cells in these mice also readily produce IL17a in response to αCD3/CD28 stimulation or the cytokine cocktail, whereas peripheral CD8αß^+^ γδ^+^ T cells from both *Foxn1*^Δ/Δ^ mice and P5 neonates are potent producers of IFNγ in response to the same stimuli (**Figures 3 and 4**).

### *Foxn1*^Δ/Δ^ mice are highly susceptible to *Bordetella pertussis* infection

*B. pertussis* is a gram-negative respiratory pathogen which causes particularly severe disease in infants. Therefore, we inoculated *Foxn1*^Δ/Δ^ mice with *Bp* to assess their susceptibility to infection compared to *Foxn1*^+/Δ^ controls. *Foxn1*^Δ/Δ^ mice harbored a significantly higher bacterial burden in the lungs at all timepoints, and in the nasal cavity at 28 dpi (**Figure 5A, B**). To determine if an altered T cell response underpins the susceptibility of *Foxn1*^Δ/Δ^ to higher *Bp* burden, we first assessed the overall frequency and number of CD3^+^ cells accumulating in the lungs of these mice versus the *Foxn1*^+/Δ^ controls at various times post *Bp* infection. Similar to the overall reduction in peripheral T cells in *Foxn1*^Δ*/*Δ^ mice [20], we also observed a reduction in the frequency and total cell numbers of CD3^+^ cells in the lung after *Bp* infection (**Figure 5D**). While CD3^+^ cell numbers increased over time in response to infection in *Foxn1^+/^*^Δ^ mice, the reduced numbers of the same cells in *Foxn1*^Δ*/*Δ^were unchanged after infection. Together, these data demonstrate that *Foxn1*^Δ*/*Δ^mice are defective in their ability to control *B. pertussis* infection.

**Figure 5.**
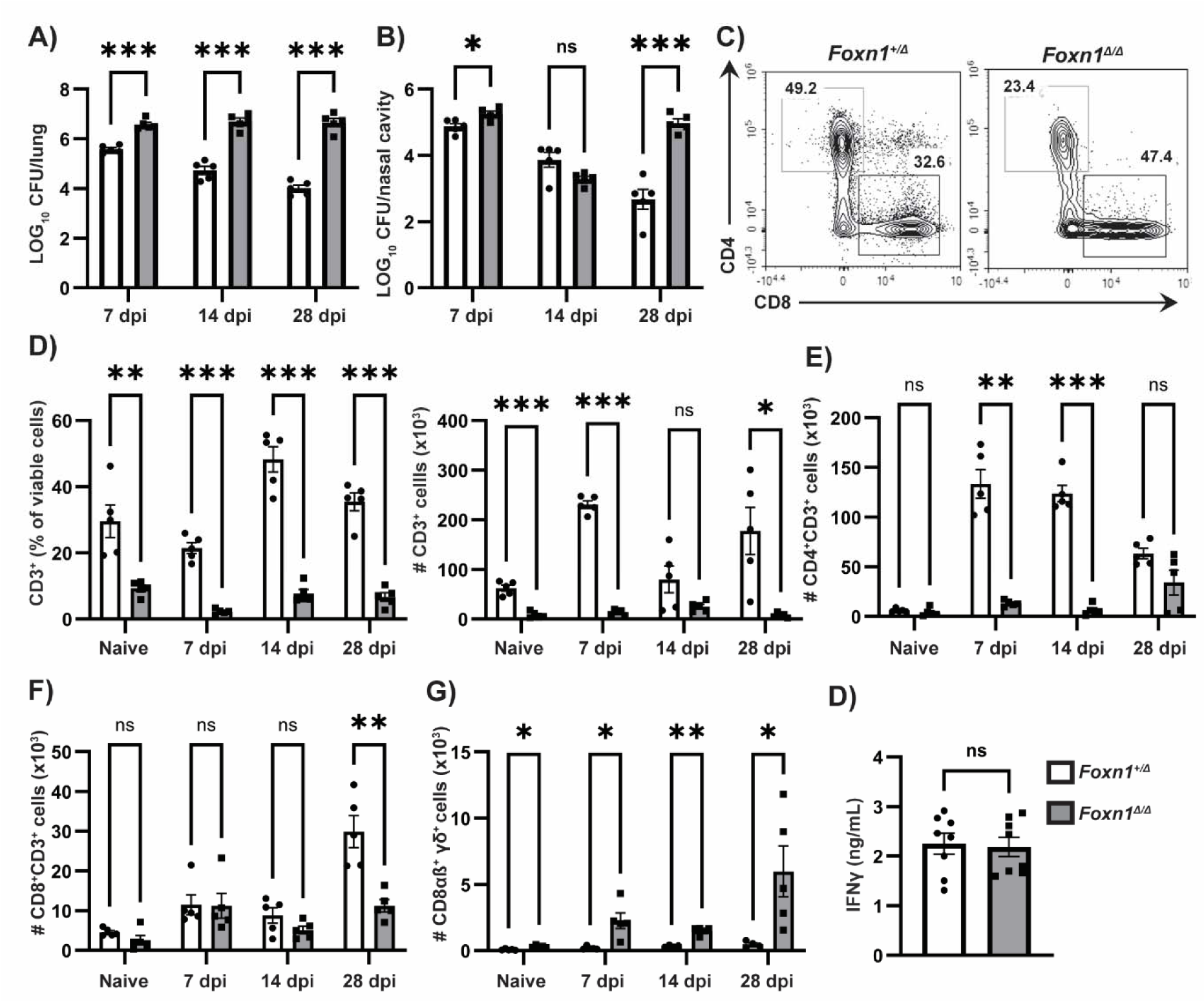
*Foxn1*^Δ*/*Δ^ mutants fail to control *Bp* infection. Bacterial burdens of *Foxn1^+/^*^Δ^ (white) and *Foxn1*^Δ*/*Δ^ (grey) isolated from the (A) lungs and (B) nasal cavity. (C) Representative gating of lung CD4^+^ and CD8^+^ T cells in *Foxn1*^Δ*/*Δ^ and *Foxn1^+/^*^Δ^ mice, previously gated on single cell lymphocytes, CD45^+^ cells, and CD3^+^ cells. (D) Quantification of CD3^+^ cells from lungs of *Foxn1^+/^*^Δ^ control and *Foxn1*^Δ*/*Δ^ mutant mice as % of viable cells, and total number of cells (per lung). (E) Total number of lung CD4^+^ (F), CD8^+^ (G) and CD8αβ^+^ γδ T cells (E). (H) IFNγ quantification of lung homogenate supernatant from *Foxn1^+/^*^Δ^ (white) *Foxn1*^Δ*/*Δ^ (grey) at 7 d post *Bp* infection. Experiments were completed in duplicate (n=4-6 mice per group). Error bars show standard error of the mean. Statistical significance was calculated via unpaired two-tailed T test. (*p=<0.05, **=p<0.01, ***=p<0.001).

To better understand how changes in specific populations of T cells responding to *Bp* might affect susceptibility, we assessed the difference in the frequency and number of CD4^+^ and CD8^+^ T cells, of which the former is required for control and development of convalescent immunity in adult C57BL/6 mice. *Foxn1*^Δ/Δ^ mice failed to accumulate significant numbers of CD4^+^ T cells in the respiratory tract compared to *Foxn1*^+/Δ^ control mice, and these numbers remained relatively static over time (**Figure 5C, E)**, akin to the overall CD3^+^ cells in these same mice (**Figure 5D)**. By contrast, *Foxn1*^+/Δ^ harbored a significant number of CD4^+^ T cells in the lung, with numbers increasing over time after infection (**Figure 5A, C, E**). *Foxn1*^Δ/Δ^ mice had similar numbers of CD8^+^ T cells as *Foxn1*^+/Δ^ controls at all timepoints tested, aside from an increase at 28 dpi (**Figure 5F**). Interestingly, unlike all other T cells quantified, *Foxn1*^Δ/Δ^ mice accumulated a significant population of CD8αβ^+^ γδ T cells compared to control mice over time post infection (**Figure 5G**). However, there was no difference in the amount of IFNγ in lung homogenate supernatant from *Foxn1*^Δ/Δ^ and *Foxn1*^+/Δ^ mice at 7 dpi (**Figure 5H)**. Thus, the increased susceptibility to *Bp* is likely attributed to a failure to generate or recruit CD4^+^ T cell effectors that cannot be compensated for by CD8αβ^+^ γδ or other innate-like T cells enriched in the lungs of *Foxn1*^Δ/Δ^ mice (**Figure 2**).

### *Foxn1*^Δ/Δ^ mice fail to recover from influenza infection

Influenza causes a viral respiratory disease, with infants < 5 years old more susceptible to serious complications requiring hospitalization [4, 39]. T cells are critical for an effective primary response to influenza infection [40], and a significant mediator of this response is IFNγ [41, 42]. Therefore, we sought to examine the ability of the T cells in *Foxn1*^Δ*/*Δ^adult mice to respond to influenza infection. To this end, we infected *Foxn1*^Δ*/*Δ^mice with the x31 strain of influenza and monitored morbidity (weight loss) and viral burden over time. *Foxn1*^Δ*/*Δ^ and *Foxn1^+/^*^Δ^ mice responded comparably to the initial infection, with relatively mild weight loss through ∼9dpi and slightly lower viral titers in *Foxn1*^Δ*/*Δ^ mice at 3 dpi (**Figure 6A, B**). However, *Foxn1*^Δ*/*Δ^mice experienced a recrudescence of weight loss around 10 dpi when control *Foxn1^+/^*^Δ^ mice have cleared the virus and regained any weight lost due to infection. Not surprisingly, lung viral titer measured at 10 and 22 dpi revealed a persistent viral load in *Foxn1*^Δ*/*Δ^lungs (**Figure 6B**). As antigen-specific CD8^+^ T cells are vital to clearance of viral infections, we assessed the development of epitope-specific CD8^+^ T cells in *Foxn1*^Δ*/*Δ^mice using the NP_366-374_ MHC class I tetramer. *Foxn1*^Δ*/*Δ^mice failed to generate a profound NP_366–374_ specific CD8 T cell response, exhibited by a 50-fold deficit of NP-tetramer^+^ cells in the lung and associated airway (as represented by BAL) of *Foxn1*^Δ*/*Δ^mice at 10 dpi (**Figure 6C, D**). Furthermore, T cells isolated from various tissues and stimulated with NP peptide showed a markedly reduced frequency of IFNγ^+^ positive cells (**Figure 6E, F**). We observed ∼50 fold less of these cells in the lungs and BAL of FoxN1^Δ/Δ^ mice isolated 10 dpi, which for the lung coincides with the difference in the number of Tet^+^ cells recovered from this site. Therefore, it is likely the lack of NP-specific CD8^+^ T cells contributed to the failure of *Foxn1*^Δ*/*Δ^mice to fully control viral titers.

**Figure 6:**
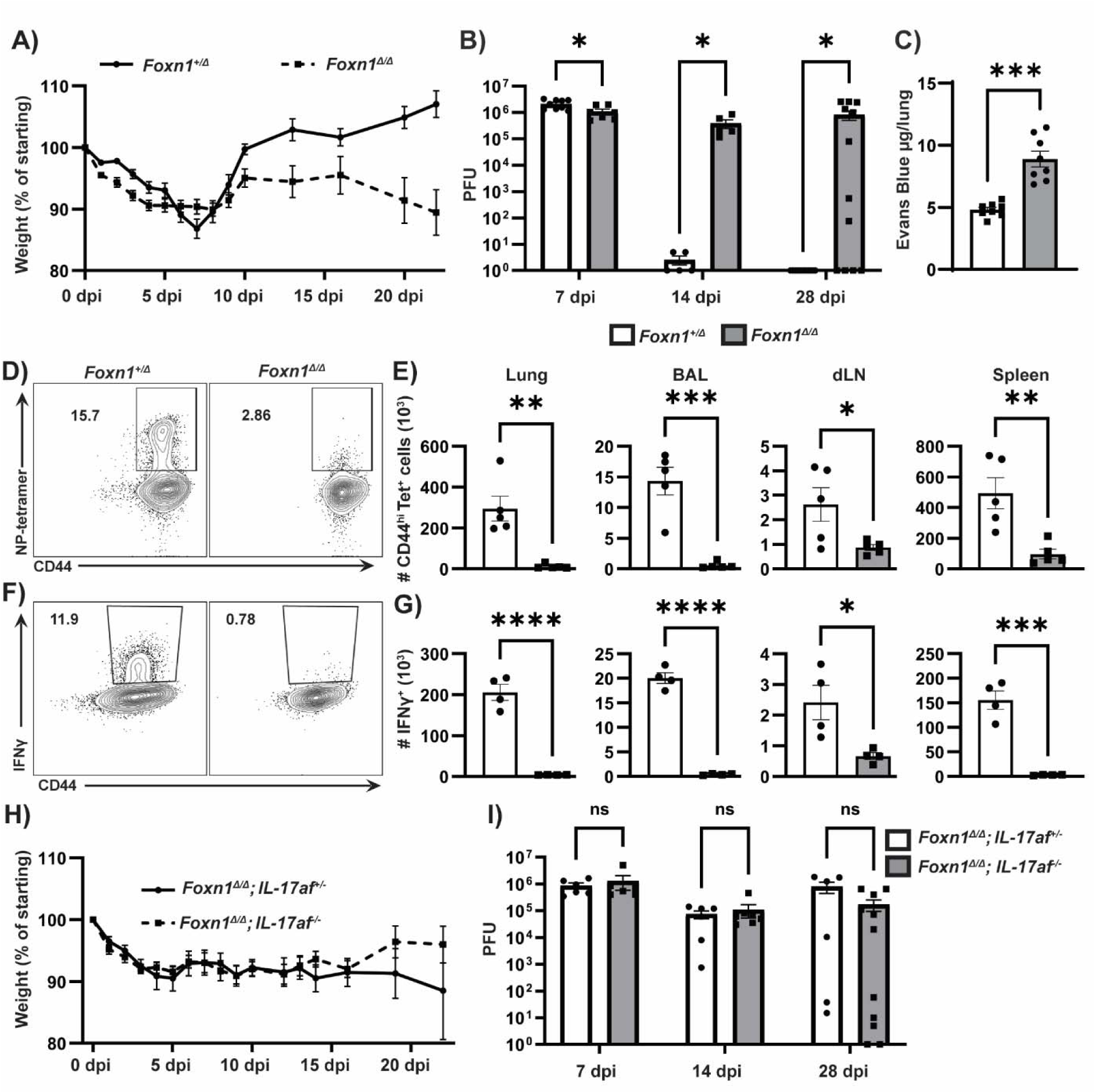
*Foxn1*^Δ*/*Δ^ mice have prolonged influenza infection. (A) Weight loss of *Foxn1^+/^*^Δ^ (white) and *Foxn1*^Δ*/*Δ^ (grey) mice following x31 infection (n=24-20 mice at day 0 reduced by 5-7 mice at 3 and 10 days post-infection). (B) Viral titer of lung homogenate supernatant from mice at the indicated time post influenza infection (n=5-12 mice per group per timepoint).(C) Quantification of Evan’s Blue dye (EB) in the perfused lung of *Foxn1^+/^*^Δ^ (white) *Foxn1*^Δ*/*Δ^ (grey) mice at 22d post x31 infection, 1 h after intravenous injection of EB. (D) Representative flow cytometry plots of NP-tetramer^+^ CD44^+^ T cells isolated from bronchoalveolar lavage (BAL); previously gated on single cell lymphocytes and CD8α^+^ (E) Quantification of the number of CD8^+^CD44^+^NP-tetramer^+^ cells in the indicated tissues and isolates at 10dpi (n=5 mice per group). (F) Representative flow cytometry plots of IFNγ^+^ CD44^+^ T cells isolated from BAL; previously gated on CD8α^+^. (G) Quantification of the number of IFNγ^+^CD8^+^ T cells after in vitro stimulation with the NP_366-374_ peptide (n=4 mice per group). (H) Weight loss of *Foxn1^+/^*^Δ^; *IL17af^-/+^* and *Foxn1*^Δ*/*Δ^; *IL17af^-/-^* mice over time following x31 infection (n=17-24 mice at day 0 reduced by 5-7 mice at 3 and 10 days post-infection). (I) Viral titer of lung homogenate supernatant from *Foxn1*^Δ*/*Δ^; *IL17af^+/-^* and *Foxn1*^Δ*/*Δ^;*IL17af^-/-^* mice at the indicated time post x31 infection (n=5-11 mice per group per timepoint). Experiments were completed in duplicate. Error bars show standard error of the mean. Statistical significance was calculated via unpaired two-tailed T test. (*p=<0.05, **=p<0.01, ***=p<0.001).

Loss of both lung vascular endothelium and alveolar epithelium barrier function is the hallmark of acute respiratory distress syndrome, a complication of respiratory infection with a high mortality rate [43]. To examine the effects of a persistent influenza infection on barrier integrity, we injected Evan’s blue dye intravenously to track leakage of serum albumin across the lung vascular endothelium into the parenchyma, and across the alveolar epithelium. Accumulation of Evan’s blue dye remaining in the perfused lungs was quantified as a measure of endothelial leakage, while BAL was collected to quantify leakage across the alveolar epithelium. While loss of vascular endothelial integrity was observed at 22 dpi, with Evan’s blue recovered from perfused lungs ∼2x higher in *Foxn1*^Δ*/*Δ^ mice compared to *Foxn1*^+/Δ^ mice (**Figure 6 B, C**). However, we did not observe significant Evan’s blue dye in the BAL of either *Foxn1*^Δ*/*Δ^ or *Foxn1^+/^*^Δ^ mice, with most samples being below the 1µg/mL limit of detection (**Figure S2.)**

Despite the fact that *Foxn1*^Δ*/*Δ^mice failed to clear the influenza infection, they appeared to limit early infection very effectively, exhibited by slightly lower viral titers at 3 dpi and mild weight loss comparable to controls. In certain settings, IL-17 can participate in anti-influenza immunity [44]. To determine if the large pool of lung embedded CD4^+^ T cells in naïve *Foxn1*^Δ*/*Δ^mice capable of producing IL-17 (**Figure 3C, D**) contributed to controlling acute influenza infection, we generated *IL-17a, IL-17f* double knockouts on the *Foxn1*^Δ*/*Δ^background (*Foxn1*^Δ*/*Δ^*; IL-17af^-/-^*). These mice were infected with x31 influenza and compared to the *Foxn1*^Δ*/*Δ^ mutant IL-17af heterozygous controls (*Foxn1*^Δ*/*Δ^*; IL-17af^+/-^*). Interestingly, the failure of *Foxn1*^Δ*/*Δ^*; IL-17af^-/-^* mice to produce IL-17 had no significant effect on weight loss (**Figure 6E**). Additionally, plaque assays from lung tissue show no significant difference in viral titer between the two groups of mice at all timepoints assessed (**Figure 6F**).

Overall, these results suggest that *Foxn1*^Δ*/*Δ^mice harbor functionally unique T cell subsets and responses, characterized by the enrichment of T cells displaying a memory and/or innate-like phenotype in barrier tissues. However, the failure of *Foxn1*^Δ*/*Δ^ mice to generate an influenza-specific CD8^+^ T cell response resulted in increased susceptibility and disease severity. Together, this work further characterizes the unique populations of T cells identified in *Foxn1*^Δ/Δ^ mice and describes the role for these cells in the increased susceptibility of *Foxn1*^Δ/Δ^ mice to both bacterial and viral pathogens.

## Discussion

Despite the fact that newborns <6 months are highly susceptible to respiratory infection [4], significant knowledge gaps remain regarding how immunity is structured early in life in non-lymphoid tissues like the lung. We and others have previously developed short-term neonatal infection models to assess early responses to various pathogens and have identified significant differences from the immune response in adults [10, 11, 25, 44–47]. While sufficient to replicate many conditions of neonatal responses, natural mouse models of neonatal infections fail to isolate factors outside of immune system development. Moreover, the immune developmental windows in mice can be brief and difficult to isolate to study age-specific responses to pathogens. Indeed, particularly in regard to T cells developing in the thymus, it is known that both the thymocyte precursor cells and thymic microenvironment are changing early in life [5, 48, 49] which may differentially affect the T cell response if it spans these timeframes. Thus, conventional murine neonatal short-term models are in part limited in their ability to study the response of the neonatal immune system, particularly during longer term infections. While neonatal thymectomies remain an effective tool, genetic disruption of thymus development is an additionally attractive strategy for producing a steady state of neonatal-like T cells.

Here, we utilized adult *Foxn1*^Δ/Δ^ mice with a persistently fetal thymus to assess T cell immunity in the lung. Given T cells generated in the early thymus are also known to have a more restricted TCR repertoire than those generated later due to the delayed expression of terminal deoxynucleotidyl transferase (TdT) [50–52], it is quite possible that the earliest T cells that exit the thymus and populate the lung have increased innate-like functions, to provide short-term protection. In our study, we observed that *Foxn1*^Δ/Δ^ mice had increased populations of tissue-embedded innate-like T cells in their lungs, while circulating T cell numbers were reduced [20]. It may be that the memory-like phenotype observed in T cells from *Foxn1*^Δ*/*Δ^ mice also includes expression of chemokine receptors and other adhesion molecules that facilitate tissue entry. Interestingly, the lungs of *Foxn1*^Δ*/*Δ^ mice were enriched for T_VM_ cells (**Figure 1C-D**), which are known to undergo proliferation in IL-15 rich environments [53]. We did observe phenotypic differences in lung T_VM_ cells from *Foxn1*^Δ/Δ^ mice compared to *Foxn1*^+/Δ^, with those from the *Foxn1*^Δ/Δ^ mice expressing high levels of CD49d (**Figure E**), an integrin that is also indicative of previous antigen exposure. This is an unusual observation given that T_VM_ are known to home to non-lymphoid tissues [54], are enriched early in life [5], and typically reported as CD49d^lo^ [55]. Interestingly, T_VM_ from the spleens of *Foxn1*^Δ/Δ^ mice express low or no CD49d (**Figure 1H**), suggesting this atypical phenotype is tissue specific. This raises some interesting possibilities: 1) that CD49d expression is indicative of an unknown encounter to commensal or self-antigens in the lung; or 2) that some thymocytes exiting the fetal thymus retain CD49d expression which facilitates entry into non-lymphoid tissues like the lung; or 3) that simply not all neonatal T_VM_ are CD49d^lo^. Future experiments will attempt to determine the origin of CD49d expression on T cells in the lungs of *Foxn1*^Δ/Δ^ mice and whether these are functionally bon-fide T_VM_.

We also observed that naïve *Foxn1*^Δ/Δ^ mice harbored a large frequency of CD8αß^+^ γδ T cells both circulating through and embedded in the lung tissue. There are relatively few studies describing these cells; however, recently published work from the Pennington and Silvo-Santos groups aligns with our findings [56]. They reported that CD8αß^+^ γδ T cells mature in the perinatal thymus, produce IFNγ, and are particularly responsive to bystander activation. Indeed, we also observe that these neonatal-like CD8αß^+^ γδ T cells express significant amounts of IFNγ when stimulated with IL-2/12/18 (**Figure 4B**); however, given that we are stimulating bulk splenocytes, we cannot rule the possibility that other cytokines produced by accessory cells in the culture activate these cell by other methods. Nonetheless, these same CD8αβ^+^ γδ T cells also express IFNγ when stimulated via the TCR (**Figure 4D**). While these CD8αß^+^ γδ T cells have been described in human models of disease [17, 56, 57], their function in neonatal immunology has yet to be determined. The enrichment of these cells in adult *Foxn*1^Δ/Δ^ mice presents an opportunity to not only further study how these cells develop, but also probe their relevance in disease models.

*Foxn1*^Δ/Δ^ mice permit the investigation of unconventional T cell responses to pathogens, as they are seemingly lacking a conventional T cell response (Figure 5–6), but retain many innate-like T cells (**Figure 1-4**) [18, 19, 21]. Indeed, *Foxn1*^Δ/Δ^ mice were significantly more susceptible to *Bp* than immunocompetent *Foxn1*^+/Δ^ controls which successfully controlled *Bp.* This is despite a relatively large pool of T cells pre-positioned within the lung parenchyma (**Figure 1A, B**) and similar levels of IFNγ in the lung at 7 dpi (**Figure 5H**), suggesting expansion of conventional CD4 T cells is required for effective control of *Bp* in mouse models.

Influenza infection of the *Foxn1*^Δ/Δ^ mice allowed us to probe immunity to viral respiratory infection and compare this to bacterial respiratory infection (*Bp).* The lack of a protective NP-specific CD8^+^ T cell response in *Foxn1*^Δ/Δ^ mice (**Figure 6C-F**) is not unexpected given wild-type mice develop most NP-specific T cell clones around 7 days post birth [52], not during the fetal period. Nonetheless, these mice provide a unique opportunity to determine how innate-like T cells provide protection to influenza infection without the subsequent activation and recruitment of antigen-specific CD8^+^ T cells. Our data shows that the enriched pools of T cells in the lung can provide immediate, but not lasting or full control of influenza infection (**Figure 6**). Given that the respiratory tract of *Foxn1*^Δ*/*Δ^ (**Figure 1**–**2**) and neonatal mice (**Figure 2D**) [10] are enriched for tissue resident and circulating T_VM_ and γδ^+^ T cells, these cells may provide an important front-line defense until conventional antigen specific T cells arrive. When the latter cells are missing, as in the *Foxn1*^Δ/Δ^ mice, infection cannot be fully controlled, resulting in increased morbidity (**Figure 6A**). This antigen-specific deficiency likely extends to T_FH_ and ultimately antibody responses as the infection remained high in these mice well past the point at which high-affinity antibodies would lead to control of the infection [58]. Thus, an adequate T cell repertoire is required for an efficient neonatal response to influenza. Future studies will prioritize characterization of the antibody response and T cell-help in the germinal center of *Foxn1*^Δ/Δ^ mice.

As mentioned previously, the innate-like T cells enriched for in the lungs of neonatal mice have been shown by our laboratory and others to be potent producers of IFNγ [17]. IFNγ expression in non-lymphoid tissues like the lung is known to induce other alarm signals to draw circulating effector and memory CD8^+^ T cells to the site of infection. However, other studies have implicated specific cytokines in early neonatal immune responses to respiratory infection. For example, the Thomas laboratory demonstrated that IL-17 producing γδ T cells play a role in controlling early influenza infection in neonates [44]. However, our data does not support a role for IL-17 in controlling acute influenza infection when thymus development is arrested in the late fetal period, due to the observation that loss of IL-17 in the *Foxn1*^Δ/Δ^ mice does not exacerbate infection or disease (**Figure 6H-I**). There are major differences in the experimental design between our studies, including but not limited to the inoculating virus, but importantly their infection of neonates at P7. This is a highly transitional period in the thymus, where growth is subsiding, TECs and the thymic microenvironment are dynamically changing [49], and the thymocyte precursor is transitioning from a fetal-derived hematopoietic HSC to a bone marrow-derived HSC [5]. Therefore, there may be a highly ordered development of T cells early in life that the HSC precursor and thymic microenvironment control over time, layering the immune response to one that is appropriate in the context of the rapidly developing organism. For these reasons, it will be important to isolate and characterize the immune potential of neonatal T cells to discrete points in time within the neonatal period to consider the role of each layer. Timed thymectomy, cellular timestamping, and other genetic models will be used in the future to better define these temporal parameters. Nonetheless, we propose that adult *Foxn1*^Δ/Δ^ mice may be a useful addition to influenza research, especially for studying innate-like T cell responses.

In summary, we characterize the unconventional T cell subtypes and functions in the lung of *Foxn1*^Δ/Δ^ mice and describe their primary response to two pathogens which disproportionately affect infants. These results demonstrate increased susceptibility of *Foxn1*^Δ/Δ^ mice to *Bp* and influenza infection (**Figure 5-6**). Interestingly, *Foxn1*^Δ/Δ^ mice recapitulated some facets of the neonatal T cell compartment such as CDαß γδ T cells and increased frequency of T_VM_. Also, the lack of an influenza specific T cell response in *Foxn1*^Δ/Δ^ mice is likely a result of low TCR diversity, a property of the neonatal T cells. However, the propensity for CD4 T cells from *Foxn1*^Δ/Δ^ mice to produce IFNγ and IL17a in response to various stimuli is not a feature of neonatal CD4 T cells (**Figure 1-2**), which are predominantly suppressive [5, 6, 9]. Also, *Foxn1*^Δ/Δ^ mice are lymphopenic and it is well known that lymphopenia induced proliferation can produce changes in T cell phenotypes. This is particularly relevant to our analysis of T_VM_ in these mice, as this phenotype can be induced by lymphopenia induced proliferation alone [59]. Lastly, while T cells from adult *Foxn1*^Δ/Δ^ mice may be comparable to T cells made in the fetal thymus, other cells that participate in adaptive immunity such as antigen presenting cells and B cells are likely identical to that of a wild type adult mouse. Thus, *Foxn1*^Δ/Δ^ mice are not a complete representation of neonatal T cell immunity but can provide insights into innate-like cells generated in the fetal thymus. Additionally, the observation of a relatively large population of CD8αß^+^ γδ T cells in these animals suggests that the *Foxn1*^Δ/Δ^ mice could be invaluable to further characterize this novel and understudied T cell subset unique to neonates. Finally, the significant proportion of T_VM_ in the lungs of *Foxn1*^Δ/Δ^ mice also warrants additional investigation, as their developmental pathways and function in neonates to pathogens and/or allergens is largely unknown. Overall, the neonatal immune responses should be considered separate from the adult, and thus vaccines and treatments should be specifically developed for the unique neonatal immune system, a feat which can may be supported by the *Foxn1*^Δ/Δ^ model.

