## Supplemental Figures 1-3 for "Adult mice with neonatal-like T cell subsets exhibit increased susceptibility to *Bordetella pertussis* and influenza infection"

A)

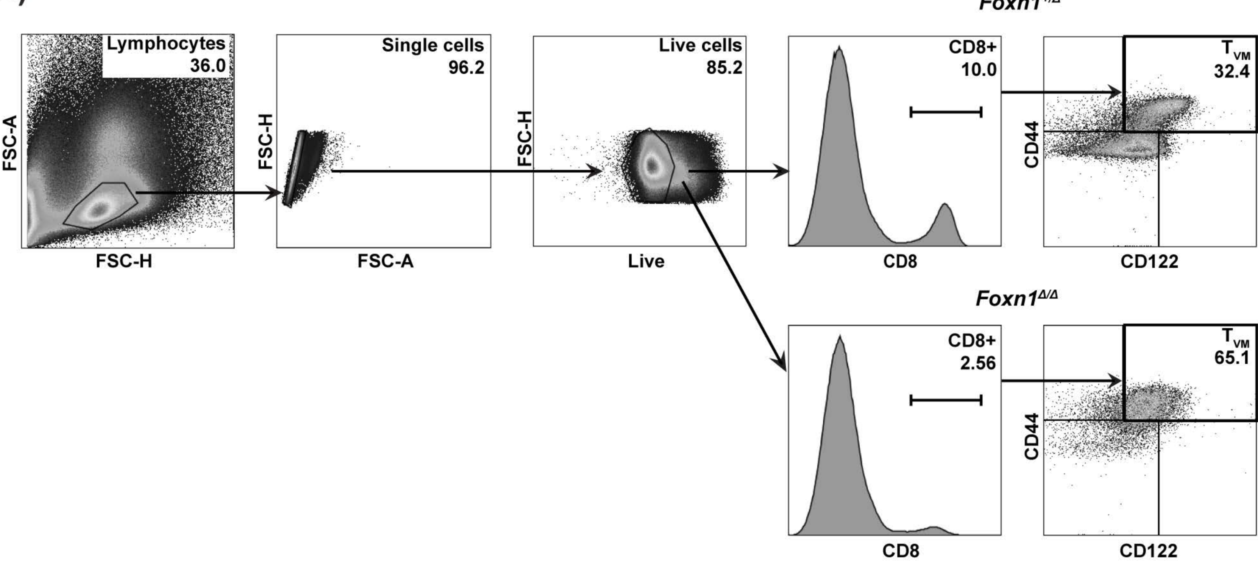


**Supplementary Figure 1: T virtual memory cell (TVM) gating strategy.** (A) Representative gating strategy of $T_{VM}$ cells; gated on lymphocytes, single cells, live cells, CD8α$^{+}$, CD44$^{+}$, and CD122$^{+}$. *Foxn1*$^{+/\Delta}$ (top) and *Foxn1*$^{\Delta/\Delta}$ (bottom).

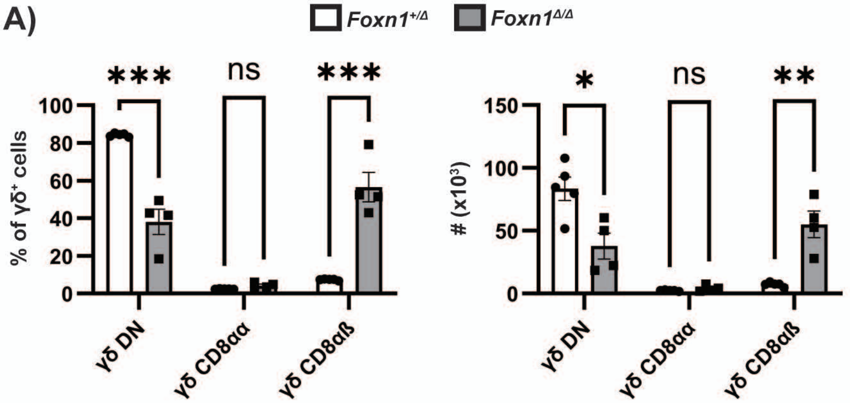


**Supplementary Figure 2: γδ T cells subset quantification in the spleen.** (A) Frequency within the γδ⁺ gate (left) and absolute number (right) of γδ T cell subsets (DN, CD8αα, and CD8αß) from *Foxn1*$^{+/\Delta}$ (white) and *Foxn1*$^{\Delta/\Delta}$ (grey) spleens.

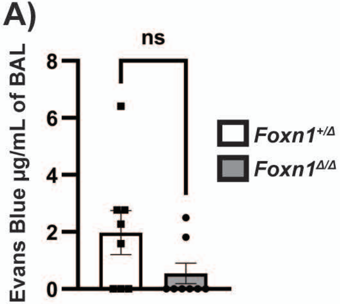


**Supplementary Figure 3: Quantification of Evan's Blue (EB) in bronchoalveolar lavage (BAL).** (A) Concentration (µg/mL) of EB recovered from the BAL of *Foxn1*$^{+/\Delta}$ (white) and *Foxn1*$^{\Delta/\Delta}$ (grey) mice 1 h after intravenous injection of EB, performed at 22 d post x31 influenza infection.
